# A SARS-CoV-2 targeted siRNA-nanoparticle therapy for COVID-19

**DOI:** 10.1101/2021.04.19.440531

**Authors:** Adi Idris, Alicia Davis, Aroon Supramaniam, Dhruba Acharya, Gabrielle Kelly, Yaman Tayyar, Nic West, Ping Zhang, Christopher L.D. McMillan, Citradewi Soemardy, Roslyn Ray, Denis O’Meally, Tristan A. Scott, Nigel A. J. McMillan, Kevin V. Morris

## Abstract

Coronavirus disease 2019 (COVID-19) is caused by severe acute respiratory syndrome coronavirus 2 (SARS-CoV-2) infection in humans. Despite several emerging vaccines, there remains no verifiable therapeutic targeted specifically to the virus. Here we present a highly effective siRNA therapeutic against SARS-CoV-2 infection using a novel lipid nanoparticle delivery system. Multiple small-interfering RNAs (siRNAs) targeting highly conserved regions of the SARS-CoV-2 virus were screened and three candidate siRNAs emerged that effectively inhibit virus by greater than 90% either alone or in combination with one another. We simultaneously developed and screened two novel lipid nanoparticle formulations for the delivery of these candidate siRNA therapeutics to the lungs, an organ that incurs immense damage during SARS-CoV-2 infection. Encapsulation of siRNAs in these LNPs followed by *in vivo* injection demonstrated robust repression of virus in the lungs and a pronounced survival advantage to the treated mice. Our LNP-siRNA approaches are scalable and can be administered upon the first sign of SARS-CoV-2 infection in humans. We suggest that an siRNA-LNP therapeutic approach could prove highly useful in treating COVID-19 disease as an adjunctive therapy to current vaccine strategies.

## Introduction

Coronaviruses have been previously linked to public health crises including the severe acute respiratory syndrome coronavirus 1 (SARS-CoV-1) outbreak in 2003 and the Middle East Respiratory Coronavirus (MERS-CoV) in 2012. These viruses led to approximately 8,096 infections for SARS-CoV-1 and 1,728 infections for MERS (WHO reports, 2004 and 2016). In contrast, the highly transmissible novel SARS-CoV-2 virus quickly escalated to a pandemic with over 128 million cases reported worldwide along with multi-organ failure, acute respiratory distress syndrome and death in the elderly and in those with underlying morbidities. The race to develop a SARS-CoV-2 vaccine began swiftly and is ongoing, however the emergence of viral variants has demonstrated the limited effectiveness of some vaccines to these variants (*1, 2*). These observations suggest an urgent and unmet need for SARS-CoV-2 specific therapies to treat coronavirus disease 2019 (COVID-19). While Dexamethasone and Remdesivir appear to provide some benefit to COVID-19 patients (*3*), a therapeutic targeted to directly inhibit SARS-CoV-2 is lacking.

RNA encodes the genome of Coronaviruses, rendering them highly susceptible to RNA interference (*4*) (*5-7*), particularly when delivered to the lungs of primates (*8*). Small interfering RNAs (siRNAs) are short double stranded RNA molecules that induce gene silencing at the transcriptional or post-transcriptional level and can be delivered to the lungs through either intranasal or intravenous routes (*9, 10*). We report here the screening of several siRNAs targeted to highly conserved regions of SARS-COV-2 that block virus expression and replication. Moreover, we find that the top candidate siRNAs are able to functionally repress virus expression *in vivo* and inhibit the emergence of COVID-19 disease when delivered intravenously using particular lipid nanoparticle (LNP) siRNA formulations.

## Results

### siRNA targeting SARS-CoV-2

To determine the effectiveness of RNAi to SARS-CoV-2 we designed several siRNAs targeted to the ultra-conserved regions in the RNA dependent RNA Polymerase (RdRp), Helicase (Hel), and 5’ untranslated leader region (5’UTR). Ultra-conserved siRNAs that target structurally accessible regions were discovered by (*a*) characterizing the 29,903bp RNA genome of SARS-CoV-2 for structural features (*11*), (*b*) sequence conservation (*12*), (*c*) RNA modifications (*13*) and (*d*) the absence of seed sequences in the human transcriptome. We used these data to prioritize approximately 9500 candidate siRNAs generated by OligoWalk (*14*) and DSIR (*15*). In addition, 163 experimentally validated SARS-CoV-1 siRNAs were assessed for homology with SARS-CoV-2 (*16*). From this stringent bioinformatic approach 18 siRNAs were selected **(Figure 1A, Table S1)**. The panel of siRNAs screened displayed varying effects on SARS-CoV-2 *in vitro* (**Figures 1B-C)** with siRNAs Hel1, Hel2, siUC7 and siUTR3 demonstrating the most potent and dose-dependent repression of virus expression (**Figure 1D**). Candidates siHel1 and siHel2 are within highly conserved regions and are able to target both SARS-CoV-1 and SARS-CoV-2 (**Figures S1A-B**).

**Figure 1.**
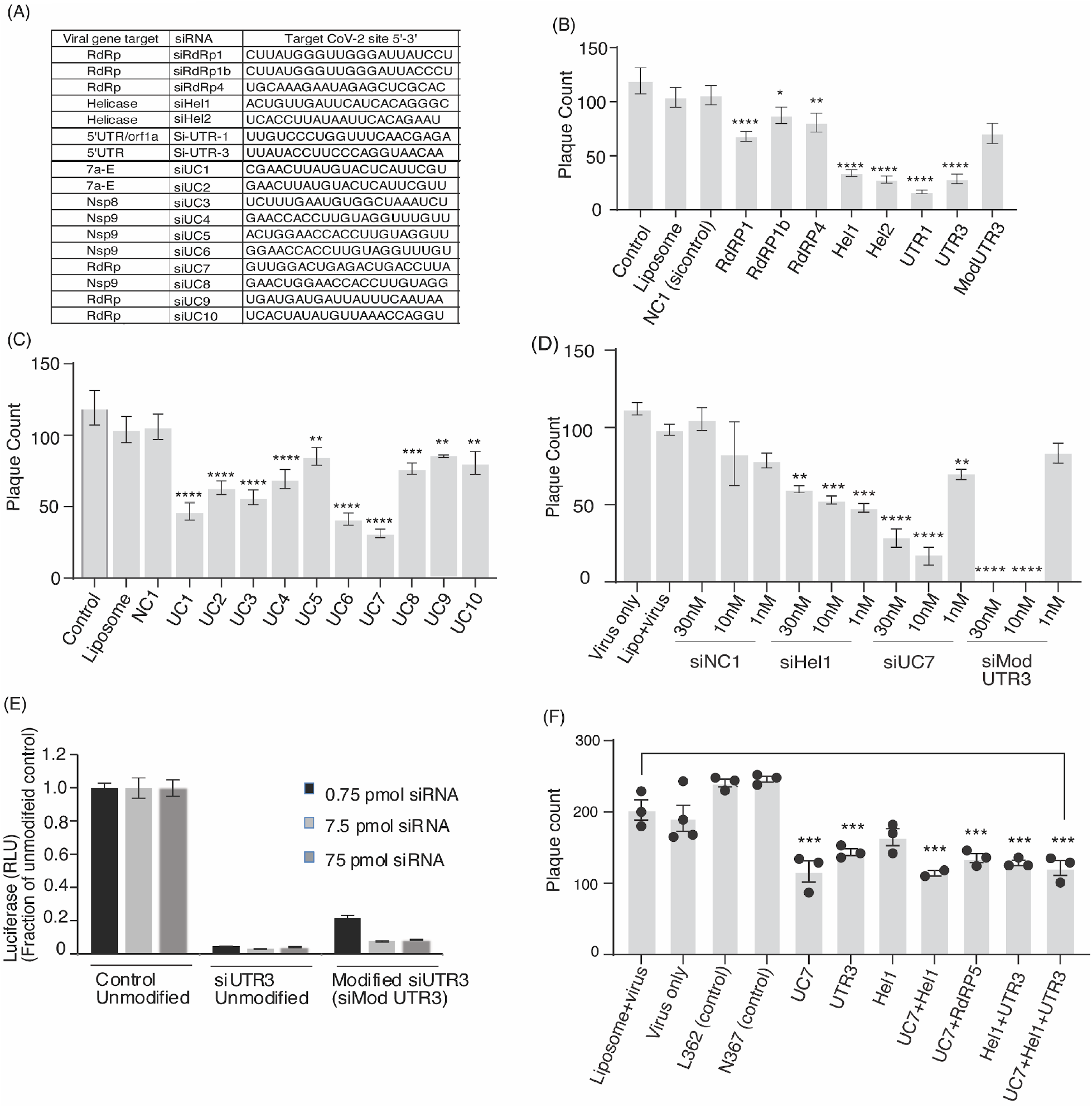
SiRNA screening against SARS-CoV-2. (**A**) The top candidate siRNAs selected for screening against SARS-CoV-2. (28) VeroE6 cells were either pretreated without (Liposome, Lipo+virus) or with 30nM of siRNA complexed with Lipofectamine 2000 for 24 hours before infection. Viral plaques were counted after 4 days. SiRNAs targeting genes (**B**) and phylogenetically conserved regions (**C**) were tested. (**D**) The top repressive siRNAs were screened for dose-dependent repression of SARS-CoV-2. (**E**) The resultant unmodified siRNA controls and the modified siMod UTR3 were transfected with a pSI-Check reporter vector with the 5’UTR cloned downstream of Renilla luciferase, and knockdown of luciferase activity of the modified siRNA determined relative to the unmodified control. The average of triplicate treated HEK293 cells is shown with the standard deviation. (**F**) Combinations of the top candidate siRNAs were selected, mixed in equal molar ratios to a final concentration of 30nM and assessed for repression of SARS-CoV-2 in vitro. For B-D and F triplicate treated cells are shown with the standard error of the mean of triplicate treatments and * represents level of statistical significance where p values of <0.05 (27), <0.01 (**), <0.001 (***) and <0.001 (****) were considered statistically significant as determined by one-way ANOVA analysis (Dunnett’s post-test) when compared against virus only (Control).

Chemical modifications can be used to stabilize siRNAs, which results in a longer-term expression and persistence *in vivo* and generally more potent repression (*17*). We selected siUTR3, as this target site resides in stem loop 1, a highly conserved region in the 5’UTR required for downstream transcriptional processing and expression of several viral RNAs (*18*). We find that 2’ O-methyl chemical modifications embedded into siUTR3 (**Figure S2**) exhibit increased stability in serum (**Figure S2**) and that repression of SARS-CoV-2 is maintained, although it is less potent than the non-modified siUTR3 (**Figures 1B and E**). We also found that none of the siRNAs tested demonstrated any observable immunostimulatory activity on human macrophages (**Figures S3A-B**).

SARS-COV-2 is able to rapidly evolve mutations that make the virus refractory to antibody targeting (*1, 2, 19*). It is well known with other RNA viruses, like human immunodeficiency virus (HIV), that single siRNA targeting results in the emergence of viral resistance (*20*) while combinations of siRNAs have been shown to hamper the emergence of resistant variants (*21*). To ascertain if combining siRNAs can functionally target SARS-CoV-2 we selected and screened three highly repressive siRNAs (siUTR3, siUC7 and siHel1) alone and in combination for repression of virus expression. Interestingly, we find that mixtures of siRNAs offered the same viral knockdown as was observed with single targeted siRNAs, even though the concentration of individual siRNAs in each combination was proportionally lower (50% for two siRNAs and 33% for three siRNAs)(**Figure 1F**). Collectively, these data suggest that siRNAs Hel1, Hel2, siUC7 and siUTR3 either alone or in combination can potently target and repress SARS-CoV-2, that siUTR3 can be chemically modified and retain function, and that these siRNAs do not induce nuclear factor-κB (NF-κB) and interferon regulatory factor (IRF) innate immune activation pathways.

### Lipid Nanoparticle *in vivo* delivery of anti-SARS-COV-2 siRNAs

Developing therapeutic strategies for viral infections based on siRNAs has so far proved challenging, with poor clinical success primarily being the result of subpar delivery. SARS-CoV-2 infection occurs predominantly in epithelial cells of the respiratory tract and results in diffuse alveolar damage (*22*). Macrophage and monocytes are also infected with SARS (*23*) and may be one source of the observed cytokine storm in COVID-19 disease (*24, 25*). We previously developed an intravenous liposome delivery platform that resulted in robust delivery of siRNAs to the lungs *in vivo* (*9, 10*). Unlike standard liposomes, these “stealth” lipid nanoparticles (sLNPs) are formulated to be stable in serum, circulate for long periods of time, and to protect siRNA payloads from nucleases. These liposomes can be formulated based on alterations of size and composition to traffic to the lung (*9, 10*).

Previously published work fully characterized the sLNPs with an average size of 190nm, polydispersity index of 0.326, zeta potential of 52.1 millivolts (mV), and 94.8% siRNA encapsulation efficiency (*9*). Additionally, this work demonstrated that sLNPs can target the lung (∼35%), liver (∼55%) and spleen (∼10%)(*9, 10*). Recent studies have found that increased concentrations of 1,2-dioleoyl-3-trimethylammonium-Propane (DOTAP) with DLin-MC3-DMA (MC3) into the LNP formulations results in enhanced targeting to the lung (*26*). As such we sought to contrast earlier formulated sLNPs containing 50% DOTAP with next generation modified LNPs containing 40% DOTAP+MC3 (dmLNP) for delivery of anti-SARS-CoV-2 siRNAs *in vivo* using the K18-hACE2 mouse model of COVID-19 disease (*27*). First, to validate the pathogenicity of SARS-CoV-2 in the K18-hACE2 mouse model, mice were inoculated with 4×10^4^ PFU of virus. Within the first 4 days dramatic weight loss of ∼20% of body weight, occurred (**Figure S4A**) with a corresponding heightened clinical score (**Figure S4B**). Infected K18-hACE2 mice also exhibited high viral load in the lung (**Figure S4C**), and brain (**Figure S4D**), which was infectious upon serial passage (**Figure S4E**). These data demonstrate, similar to previous observations with SARS-CoV-1 (20, 23), that the K18-hACE2 mouse model exhibits COVID-19 disease when infected with SARS-CoV-2.

Next, to determine the ability of the sLNP-siRNAs to functionally repress SARS-CoV-2, K18-hACE2 mice were inoculated with 1×10^4^ PFU of virus and treated with various sLNP-siRNA formulations intravenously (**Figure 2A**). We find that sLNP-siRNA treatment provided a survival advantage in the sLNP-siUC7 and sLNP-siHel2 treated mice compared to virus infected and sLNP-siRNA control treated mice (**Figure 2B**). The functionally treated mice exhibited less weight loss (**Figure 2C**) and an overall lower clinical score (**Figure 2D**) when compared to the control sLNP-siRNA and virus infected mice. Notably, both the sLNP-siUC7 and sLNP-siHel2 treated mice functionally repressed SARS-CoV-2 *in vivo* at day 3 based on viral outgrowth analysis from lung (**Figure 2D)** but this effect was lost by day 6, suggesting that the repressive effect of the siRNAs is transient and found ∼24-48hrs following sLNP-siRNA treatment. Markedly, the siRNA treated mice appeared most closely aligned with the mock treated mice when the transcriptomic profile of lung from day 6 was characterized (**Figure 2F and Table S2**). The sLNP-siRNAs assessed here contain ∼50% DOTAP which is a cationic lipid often used in nanoparticle and liposome formulations. Cationic liposomes can aggregate and lead to accumulation in the spleen, liver and lung. Notably, the sLNP-siRNA formulations did not appear to exhibit any overt splenomegaly in treated virus infected mice as determined by post-mortem assessment of the spleens from treated and control animals at day 3 (**Figure S5A**) and day 6 (**Figure S5B**) post-treatment. Collectively, these data demonstrate that intravenous (IV) injected sLNP-siRNAs can repress SARS-CoV-2 *in vivo* and delay the onset of COVID-19 symptoms and that siHel2 appears to be a potent siRNA for repressing viral expression when delivered intravenously with sLNPs.

**Figure 2.**
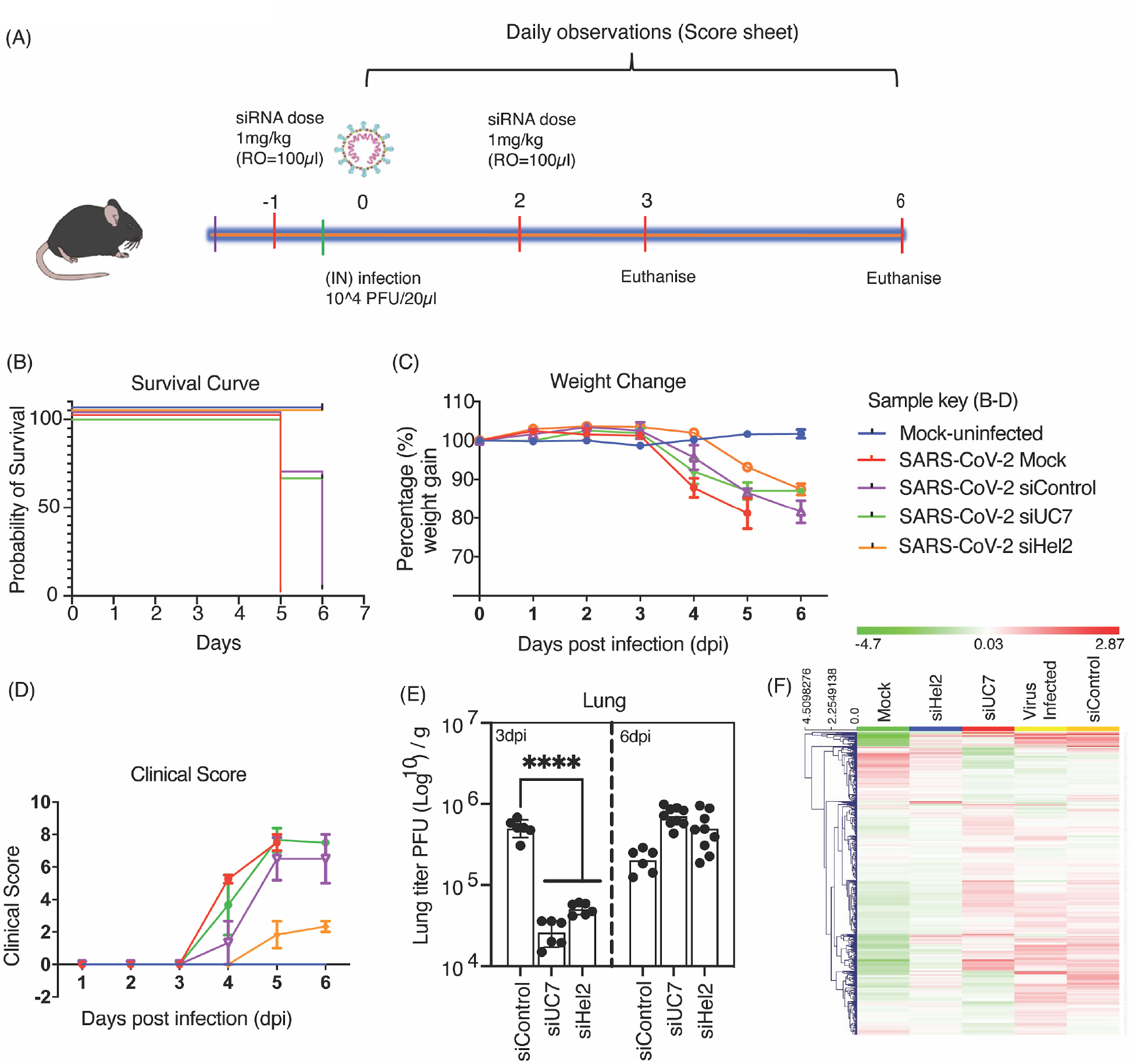
Intravenous administered sLNP-siRNA repression of COVID-19 in vivo. (**A**) 7–13-week-old K18-hACE.2 female and male mice were intranasally infected with either PBS or 10^4^ PFU/ 20 µL of SARS-CoV-2 (Australian VIC1 strain, passage 4). (**A**) Mice were intravenously treated with 1 mg/kg in 100 µL of siRNA packaged into HFDM lipid nanoparticles (LNP) by retro-orbital administration at -1 and 2 days post infection (dpi). At 3dpi and 6dpi lung and brain tissues were harvested and homogenized for immunoplaque assays. (**B-D**) Mouse survivorship, probability of survival, body weight (weight change), and clinical score were evaluated at the indicated days post-infection (dpi). (**B**) Weight loss >15% was taken as an endpoint and mice were euthanized. (**C**) Mice were weighed and scored daily until the experimental end point (6 dpi), for disease progression. (**D**) The clinical score was evaluated based on locomotion, behaviour and appearance. Each data point represents the average ± SEM of 3 to 4 mice. (**E**) The amount of infectious virus particles in lung tissues at 3 and 6 dpi were titrated by immunoplaque assays on Vero E6 cells, using a SARS-CoV-2 N protein specific antibody and expressed as PFU per gram of tissue. Each data point represents a technical replicate, where one mouse is equivalent to 3 technical replicates and bars represent the average ± SEM. (**F**) An unsupervised hierarchical cluster heatmap of immune gene expression in the lungs at 6dpi. Each row is a gene and each column is a treatment group. Rows are Z-score normalised (Green: low expression and red: high expression). A p value of <0.0001 (****) is considered statistically significant when assessed by 2-way ANOVA (Dunnett’s post-test) when compared against siControl.

The sLNP-siRNAs (**Figure 2**) contain 50% DOTAP which contributes to the highly positive surface charge which has been suggested to activate the immune system (*28*). To assess the immune stimulatory properties of the sLNP-siRNAs, we assessed mouse lung gene expression profiles using NanoString immune gene expression profiling analysis at day 6. Stimulation of interferon-regulated immune genes was observed between mock and virus infected (**Figure S6A**) and similar patterns of immune gene activation were observed between the virus infected and the various sLNP-siRNA treated mice (**Figures S6B-D, Table S3**), suggesting that siRNA treatment in these mice was not overtly immune-stimulatory.

While we did not observe any notable unique immune dysregulation with the sLNP-siRNAs, *in vivo*, and the siRNAs alone did not demonstrate any observable immune stimulation *in vitro* (**Figure S3**), recent work has suggested that reducing DOTAP can ameliorate LNP-siRNA immune stimulation. Based on these concerns, we screened a panel of formulations with reduced DOTAP at 40, 35, and 30% (**Figure S7A**). Our goal was to develop a next generation ‘stealth LNP’ formulation with reduced DOTAP and in turn incorporate the cationic ionizable lipid MC3 to help facilitate the endosomal release of siRNAs. We observe that our reduced DOTAP LNP formulations range from 80 to 115 nm in size and display low polydispersity values (**Figure S7B**). The zeta potential of our reduced DOTAP LNP formulations range from 17 to 23 mV (**Figure S7C**) and there was little observable difference in the zeta potential values between these new formulations despite the stepwise reduction in DOTAP (**Figure S7C**). Because highly positive surface charges of nanoparticles and liposomes are linked to toxicity (*28*) we view our reduction in surface charge as compared to the previous sLNP-siRNA formulation as a favourable step toward reducing potential toxicity. Furthermore, all formulations in our panel had ≥92% encapsulation efficiency of siRNA cargo (**Figure S7D**) and transmission electron microscope imaging of the DOTAP40 LNPs exhibited a uniform spherical shape (**Figure S7E**). Remarkably, the DOTAP40 and DOTAP40C LNPs remain stable for at least 9 months when stored at 4°C with nearly 100% retention of encapsulated siRNA (**Figure S8A-B**) and DOTAP40C LNPs retain function after six days at room temperature. Furthermore, the siRNAs are largely resistant to enzymatic degradation when encapsulated in these LNPs (**Figure S8C-D**). Based on these observations we selected the “DOTAP40C” formulation which contains the highest proportion of MC3 while also retaining a high proportion of DOTAP which is important for achieving lung delivery of the candidate SARS-CoV-2 siRNAs. To determine the ability of these newly formulated DOTAP/MP3 LNP-siRNAs (dmLNP-siRNAs, formerly identified as DOTAP40C) to effectively target the lung, DiD labelled dmLNP-siRNA formulations (**Figures 3A**) were generated and found to be ∼80 nm (**Figure 3B**) with a zeta potential of ∼18.58 mV and to encapsulate ≥97% of the control siRNAs (**Figure 3C**). When injected intravenously into mice and assessed 24 hrs later there was localization of DiD fluorescence in the lung (21%), liver (67%) and less so in the spleen (12%) (**Figures 3D-E**), which was similar to previous observations with sLNPs (*9, 10*).

**Figure 3.**
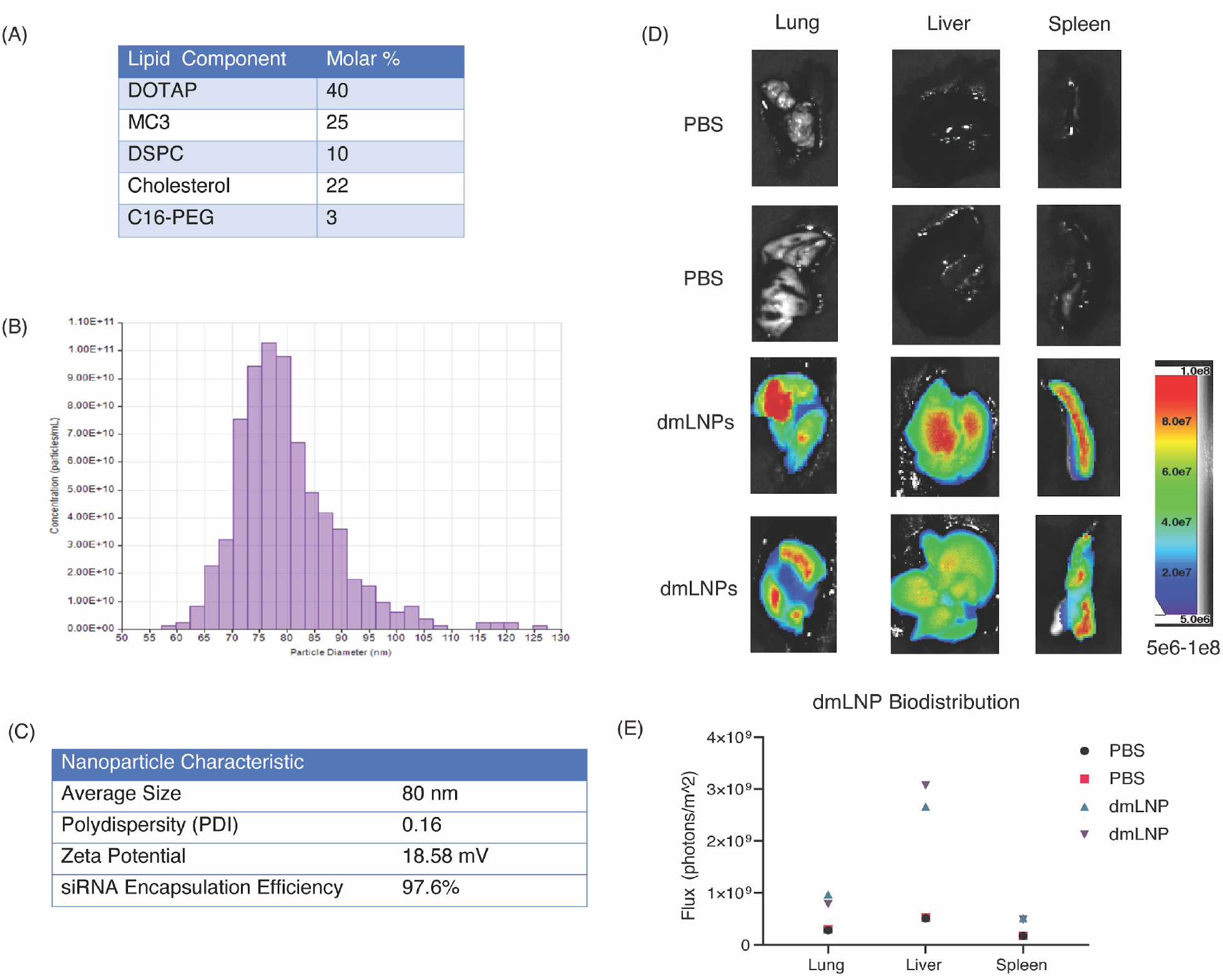
dmLNP-siRNAs characterization and biodistribution. (**A**) Molar composition of dmLNP-siRNA lipid nanoparticles. (**B**) Nanoparticle size distribution of dmLNP-siRNA lipid nanoparticles determined using the qNano Gold tunable resistive pulse sensing device. (**C**) dmLNP-siRNAnanoparticle characteristics including size, polydispersity (PDI), surface charge (zeta potential), and siRNA encapsulation efficiency. (**D**) dmLNP-siRNA biodistribution was determined in C57/BL6 mice that received DiD-labeled dmLNP-siRNA nanoparticles at 1mg/kg siRNA dose or PBS vehicle control via retro-orbital (RO) route. 24 hours after injection, mice were euthanized and the lung, liver, and spleen were removed. Organs were imaged for DiD fluorescence using a LagoX small animal imaging machine at excitation and emission wavelength of 640 and 690 nm, respectively. (**E**) Quantitative analysis of DiD fluorescence in each organ.

Next to determine the ability of the dmLNP-siRNAs to deliver functionally repressive siRNAs, K18-hACE2 mice were inoculated with 1×10^4^ PFU of virus and treated with dmLNP-siRNA formulations (**Figure 4A**). We find that mice treated with dmLNP-siHel2 exhibited a survival advantage (**Figure 4B**), exhibited less weight-loss (**Figure 4C**) and a lower clinical score (**Figure 4D**) when compared to the control dmLNP-siRNA treated and virus infected mice. Further, the data showed a recovery of weight, and a concomitant decrease in clinical score at days 6-8 in mice treated with dmLNP-siHel2 and dmLNP-siUC7, suggesting that these treatments may alleviate severe disease symptoms. Similar to previous observations with sLNP-siRNA formulations (**Figure 2E**) both the dmLNP-siUTR3 and dmLNP-siHel2 treated mice functionally repressed SARS-CoV-2 *in vivo* at day 7-8 based on viral outgrowth analysis from lung (**Figure 4E)** suggesting a *bona fide* repression of virus *in vivo*. Collectively, these data demonstrate that intravenously injected dmLNP-siRNAs, similar to sLNP-siRNA treatments (**Figure 2**), repress SARS-CoV-2 *in vivo* and delay the onset of COVID-19 symptoms and that siUC7, siUTR3 and siHel2 appear to be potent siRNAs for repressing viral expression.

**Figure 4.**
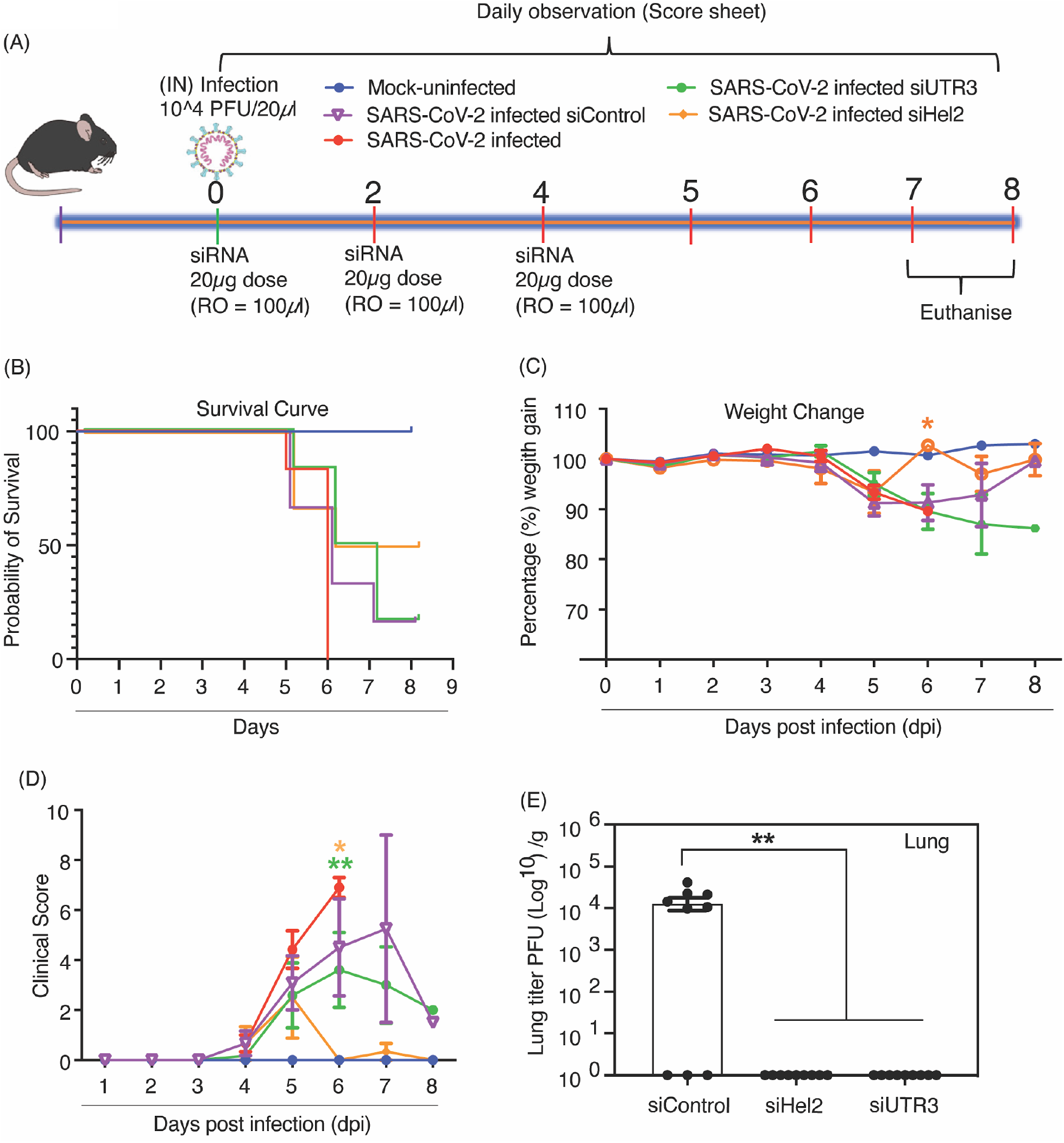
Intravenous administered dmLNP-siRNA suppression of COVID-19 in vivo. (**A**) 7– 13-week-old K18-hACE2 female and male mice were intranasally infected with either PBS or 1×10^4^ PFU/ 20 µL of SARS-CoV-2 (Australian VIC1 strain, passage 4). (**A**) Mice were intravenously treated with 1 mg/kg in 100 µL of siRNA packaged into DOTAP 40 lipid nanoparticles by retro-orbital administration at 0, 2 and 4dpi. At 6 dpi lung and brain tissues were harvested and homogenized for immunoplaque assays. (**B-D**) Mouse survivorship during infection and dmLNP-siRNA treatment, (**B**) probability of survival, (**C**) Body weight (weight change), and clinical score were evaluated at the indicated dpi. Mice which lost >15% of their initial body weight were humanely euthanized and plotted as a non-survivor. (**B-D**) Mice were weighed and scored daily until the experimental endpoint for disease progression. (**D**) The clinical score was evaluated based on locomotion, behaviour and appearance. Each data point represents the average ± SEM of 3 to 4 mice. (**E**) The amount of infectious virus particles in lung tissues at 7 and 8 dpi were titrated determined by immunoplaque assays on Vero E6 cells, using a SARS-CoV-2 N protein specific antibody and, expressed as PFU per gram of tissue. Each data point represents a technical replicate, where one mouse is equivalent to 3 technical replicates and bars represent the average ± SEM. p values of <0.05 (*) and <0.01 (**) are considered statistically significant when assessed by 2-way ANOVA (Dunnett’s post-test) against (33) SARS-CoV-2 infected only mice and (**E**) siControl.

## Discussion

Currently there are scant antivirals reported which directly target SARS-CoV-2 RNA genome. Clustered Regularly Interspaced Short Palindromic Repeats (CRISPR) has recently been used to target SARS-CoV-2 (*29*), but pre-existing antibodies to CRISPRs (*30*) and the need to translate the packaged CRISPR mRNA and gRNA in virus infected cells will hinder the clinical translation of this approach. RNAi does not require translation of mRNA, is programmable, scalable, stable and has been observed to potently repress Coronaviruses (*5-7*). We show here that RNAi and particular siRNAs, siHel1, siHel2, UC7 and siUTR3 significantly repress SARS-COV-2 *in vitro* and *in vivo* and could prove to be a useful therapeutic to treating COVID-19 disease. However, delivery of siRNAs to sites of disease, such as the lungs in COVID-19 afflicted individuals, has remained enigmatic.

The persistent cough and shortness of breath endemic in COVID-19 disease highlight the lungs as a site of significant stress and inflammation during SARS-CoV-2 infection. The thick mucosa associated with COVID-19 will likely impede the delivery of aerosolized therapeutics to infected tissues and additionally, nebulizers increase droplet dispersion which could lead to infectious particles remaining in the air thereby increasing the risk of the disease spreading. For these reasons we surmised that an intravenous route of administration as a backdoor delivery system, might prove both safe and effective. Building on this notion we turned to “stealth” LNPs (*9, 10*), which have been shown to deliver siRNAs to the lung, liver and spleen following an intravenous administration.

Recent work by Cheng et al., 2020 demonstrated that LNP formulations can be tuned to specifically target the lung by adjusting the amount of DOTAP incorporated into the particles (22). Increasing the DOTAP concentration >50% has been reported to result in lung specific expression of a luciferase mRNA reporter (*26*). Notably, this group used a combination of DOTAP (50%) and MC3 (25%) to achieve efficient LNP lung targeting. Our sLNP-siRNAs containing 50% DOTAP and no MC3 target the lung, however we find the liver and spleen are also targeted (*9, 10*). Our dmLNP-siRNAs contain 40% DOTAP and 25% MC3, but also display targeting of the lung, liver and spleen. Notably, the dmLNP-siRNAs (**Figures 3 and 4**) contained 40% DOTAP and exhibited a concomitant reduction in lung targeting when administered intravenously, regardless of the MC3 incorporation, suggesting as others have observed, that DOTAP is the key component to targeting the lung with LNPs (*26*). DOTAP is a cationic lipid that contributes to the positive surface charge of liposomes and LNPs, and has been shown to activate the immune system resulting in systemic toxicity (*31*). By reducing the amount of DOTAP in our dmLNP-siRNAs we have reduced the positive surface charge on the particles by approximately half compared to the sLNP-siRNAs. We too observed some level of immune stimulation that was most likely the result of viral infection, and not attributed to the LNP-siRNA formulations as virus infected controls and LNP-siRNA particle treated mice demonstrated a similar gene expression profile (**Figure S6, Tables S2-3**). Notably, the siRNA repression of SARS-CoV-2 appeared to be transient, and the effect of viral suppression was lost after ∼48hrs, suggesting that the LNP-siRNA formulations are rapidly cleared from the system and that as a therapeutic, a daily intravenous regimen will most likely be required during peak viral infection. SiRNA modification represents one approach that both reduces immune stimulation and increases the half-life of the siRNAs *in vivo*. Our preliminary data suggest that minimal 2’OMethyl and phosphorothioate modifications are sufficient to increase the stability of the tested siRNA (UTR3) *in vitro*. Further optimisation of such modifications will most likely provide a highly potent and stable siRNA for *in vivo* delivery.

The SARS-CoV-2 vaccine race led by Pfizer and Moderna has opened the door for future LNP based therapies. Both Pfizer and Moderna vaccines (BNT162b2 and mRNA-1273, respectively) contain an mRNA encoding the Spike protein encapsulated in an LNP delivery vehicle. Prior to the pandemic, the only FDA approved LNP based therapy was the siRNA-LNP drug Patisiran (Onpattro) used for the treatment of polyneuropathy caused by hereditary transthyretin-mediated amyloidosis (*32*). Recent successes in the clinical translation of LNPs portend a new era in nanomedicine, whereby LNPs are now viewed favourably as *bona fide* and safe delivery vehicles for mRNAs and RNAi. Building on this realized consensus of interpretation we show here that intravenously administered stealth LNPs can deliver siRNAs as a therapeutic to treat COVID-19. While both RNAi and LNP technologies are relatively new, it is becoming evident that this next generation technology is programmable, scalable, stable, relatively safe, and has much to offer as a therapeutic to specifically target SARS-CoV-2 and treat COVID-19 disease.

## Acknowledgments

We thank Mr Hamish McMath for assistance with the BL3 facility and vivarium at Griffith University. We thank Olga Villamizar, Ryan Urak and Liliana Echavarria for their assistance with the LNP characterization and animal work at City of Hope, Duarte CA. We also thank Dr. Zhuo Li and Ricardo Zerda at City of Hope Electron Microscopy Core Facility for electron microscopy assistance.

## Author contributions

A.I., A.S., designed and conducted animal experiments, G.D., D.A., T.S., R.R., A.D., C.S. carried out experiments, D.O., R.R., T.S. designed and characterised the various siRNAs, A.D. designed, optimized, and generated dmLNP-siRNAs, Y.T., generated sLNP-siRNA formulations, N.M., K.V.M, conceived, designed the experiments and wrote the manuscript. All authors have reviewed and edited the manuscript.

