## Supplementary material for "A SARS-CoV-2 targeted siRNA-nanoparticle therapy for COVID-19": Supplmental information

**Supplemental Figures**

Figure S1

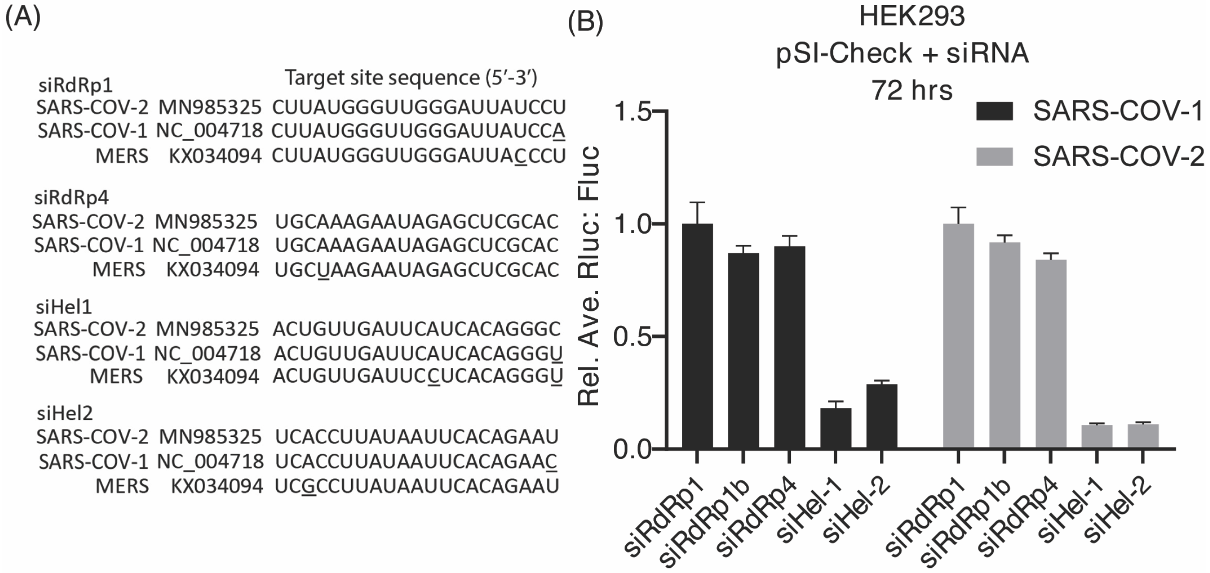


**Figure S1** *SiRNA-mediated knockdown of conserved beta-coronavirus sites in the helicase ORF. (**A**) siRNA conservation between targeting various beta coronaviruses. Alignments are shown between SARS-CoV-1, SARS-CoV-2 and MERs. Underlined DNA bases indicate a different nucleotide compared to SARS-CoV-2. **(B)** A reporter vector containing the RdRp or helicase ORF for SARS-CoV-1 and SARS-CoV-2 inserted downstream of Renilla luciferase (Rluc) was transfected with siRNAs targeting RdRp (siRdRp1,1b and 4) and helicase (siHel-1 and 2). The levels of RLuc activity was assessed at 72 hrs post-transfection, normalized to background firefly luciferase (Fluc) levels and made relative to the siRdRp1 siRNA, which was used as a negative control and set at 100%. The error bars represent standard deviation from transfections performed in triplicate.*

Figure S2

A

5’-rAmUmArCmCmUrUrCrCrCmCmArGrGrUrAmAmCrA*rA*mArCrCrAAC-3’
5’- rG*rUrUrGrGrUrUrUrGrUmUmAmCrCrUrGrGrGrArArGrGmUrAmUmArA-3’

m = 2’OMe RNA
r = RNA
* = PS linkage

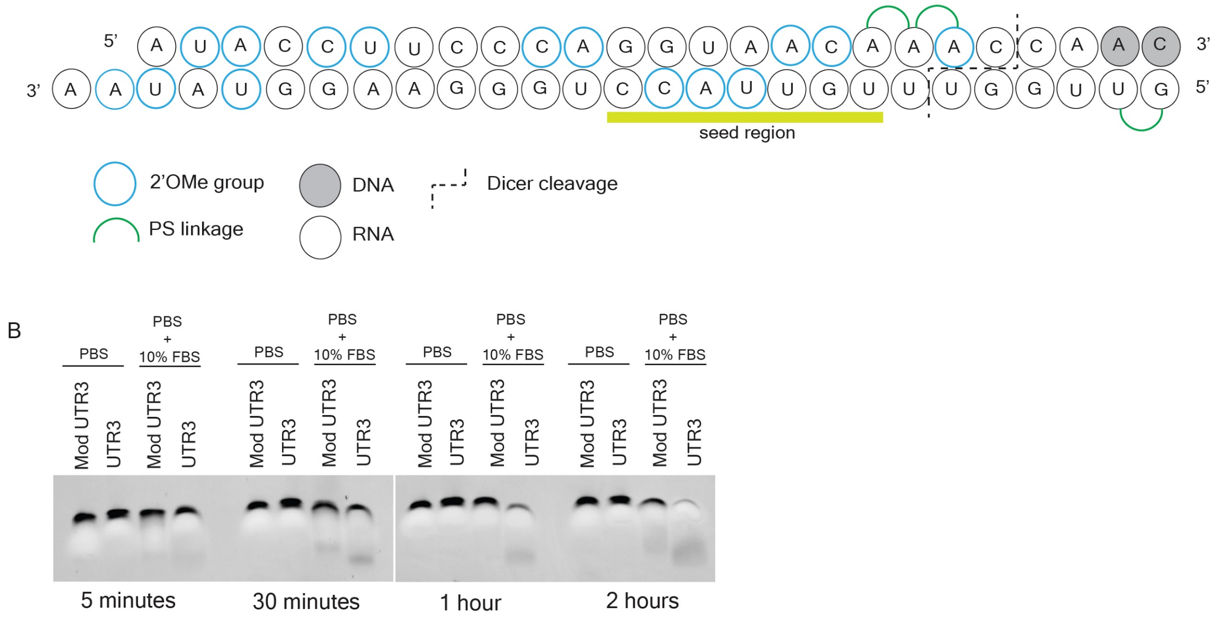


**Figure S2** *Chemical modifications imbued in dsiUTR3. (**A**) The dsiRNA UTR3 (siMod UTR3) is shown with the chemical modifications utilized to stabilize the dsiRNA and (**B**) enhance persistence while inhibiting immunogenicity. Modifications were primarily located at CA or CU regions, as these are most sensitive to nuclease activity(1). Phosphorothioate bonds were used to prevent nuclease activity and to decrease thermostability. This was preferentially done on the passenger strand at positions 19 and 20, to decrease thermostability at the 3’ end and thus promote loading of the antisense strand into the RISC complex (2). (**B**) The serum stability assay indicates the modified dsiRNA is substantially protected from nuclease attack compared to its unmodified counterpart. The assay was performed with 10 µM modified and unmodified dsiRNA in 1 X PBS or 1X PBS supplemented with 10% FBS (not heat inactivated). Samples were incubated at 37℃ for 5 min, 0.5, 1 and 2 hr time points and electrophoresed on a 6% TBE polyacrylamide gel (Novex, Invitrogen) for 30 min at 180V. Samples were stained with 2 µg/mL EtBr and images acquired under 254 nm using an EZ Imager (Bio-Rad).*

Figure S3

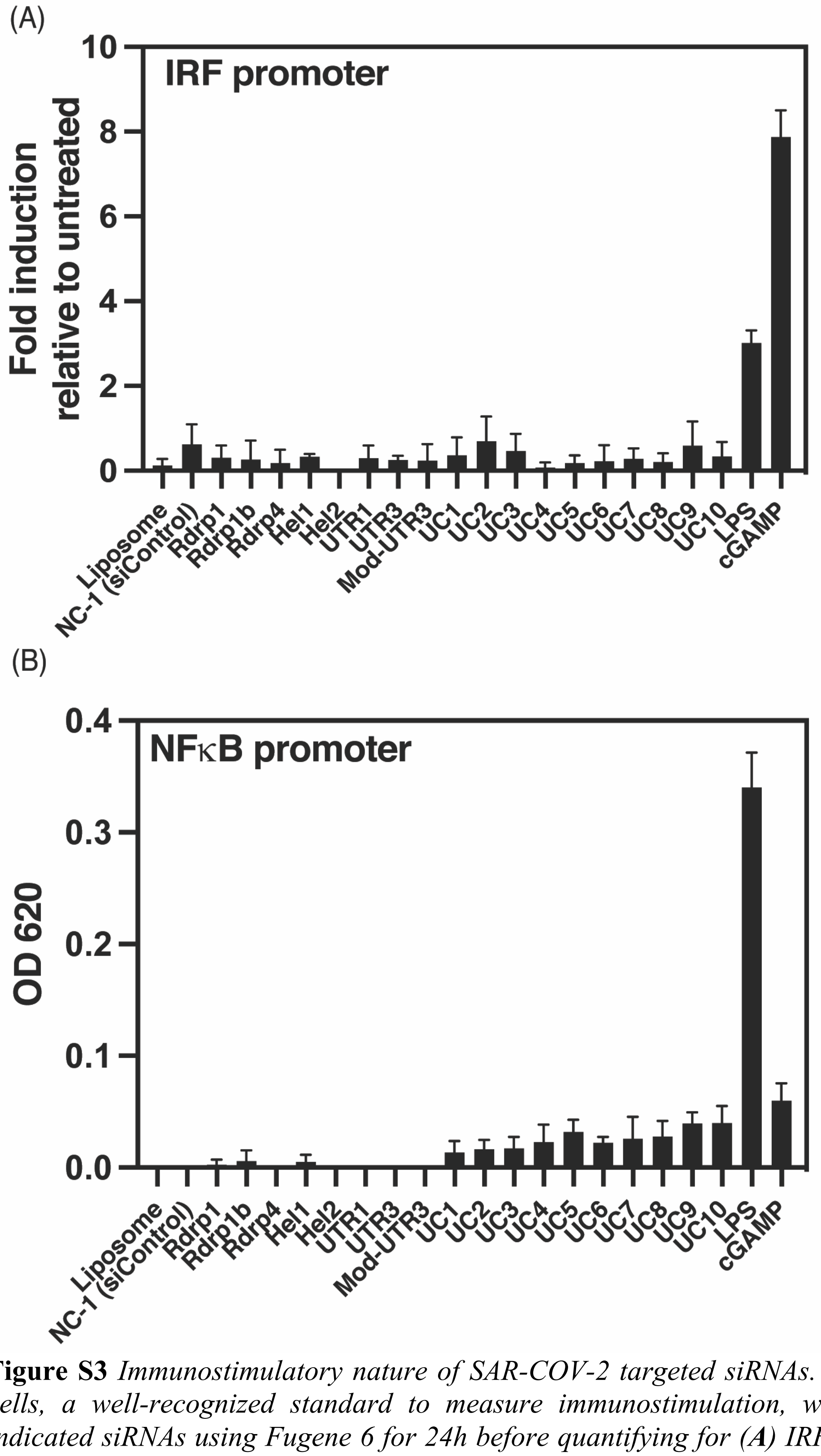


**Figure S3** *Immunostimulatory nature of SAR-COV-2 targeted siRNAs. (**A-B**) THP-1 DUAL cells, a well-recognized standard to measure immunostimulation, were transfected with indicated siRNAs using Fugene 6 for 24h before quantifying for (**A**) IRF and (**B**) NFkB gene reporter expression. Error bars denote SEM of triplicate treatments. 2'3'-cGAMP (20µg/ml) and LPS (100ng/ml) were used as positive controls for IRF and NFkB pathway stimulation, respectively.*

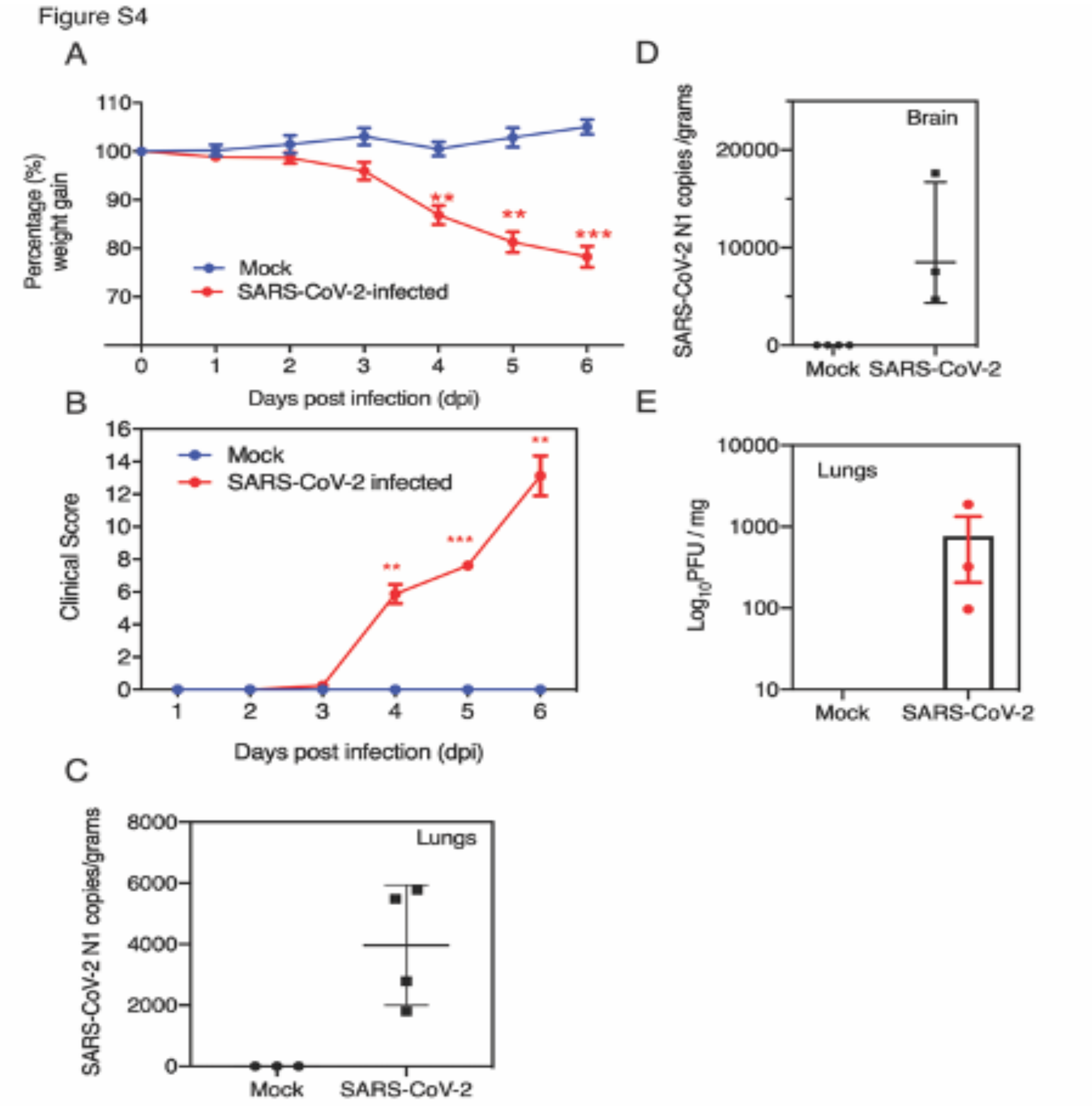


**Figure S4** *SARS-CoV-2 infection resulted in severe clinical disease and weight loss in vivo. K18-hACE2 female and male mice, 7–13-week-old, were intranasally infected with either PBS (3) or 4 x $10^4$ PFU/ 20 µL of SARS-CoV-2 (Australian VIC1 strain, passage 4). (**A-B**) Mice were weighed and scored daily until the experimental endpoint, for disease progression. (**B**) The clinical score was evaluated based on locomotion, behaviour and appearance. (**C-E**) At 6dpi lung and brain tissues were harvested and homogenized for immunoplaque assays and RNA extracted for viral copy number determination. (4) Viral copy numbers in lung and brain tissues were determined by digital droplet PCR against the N gene and expressed as viral copies per gram of tissue. (**E**) Infectious viral load in lung tissues were determined by immunoplaque assays on Vero E6 cells, using a SARS-CoV-2 N protein specific antibody and expressed as PFU per gram of tissue. (**A-B**) Each data point represents the average ± SEM of 3 to 4 mice. (**C-E**) Each data point represents an individual mouse and bars or lines represent the average ± SEM. (**A-E**) P values of <0.05 (5), <0.01 (**) and <0.001 (***) are considered statistically significant when assessed by Student t- test against mock control.*

Figure S5

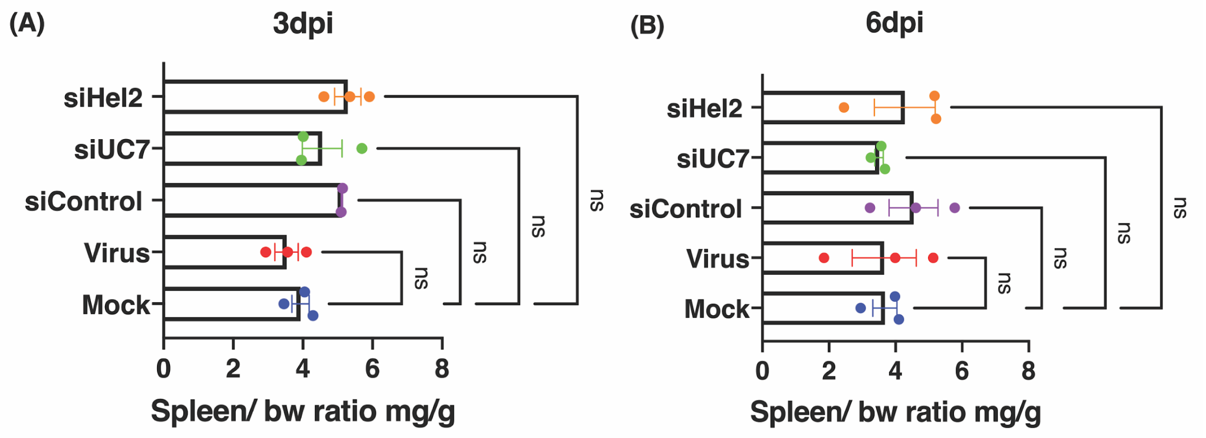


**Figure S5** *Retro-orbitally administered LNP-siRNA treatment maintained normal spleen to body weight ratios in treated mice. K18-hACE2 female and male mice, 7–13-week-old, were intranasally infected with either PBS (3) or 1x10$^4$ PFU/ 20 µL of SARS-CoV-2 (Australian VIC1 strain, passage 4). Mice were retro-orbitally (RO) treated with 1mg/kg in 100 µL of siRNA packaged into HFDM lipid nanoparticles (LNP) at days -1 and 2 post infection. (**A-B**) Mice were sacrificed at (**A**) 3 dpi and (**B**) 6 dpi and relative organ/body weight ratios of spleen was graphed. Each data point represents an individual mouse and bars represent the average ± SEM.*

Figure S6

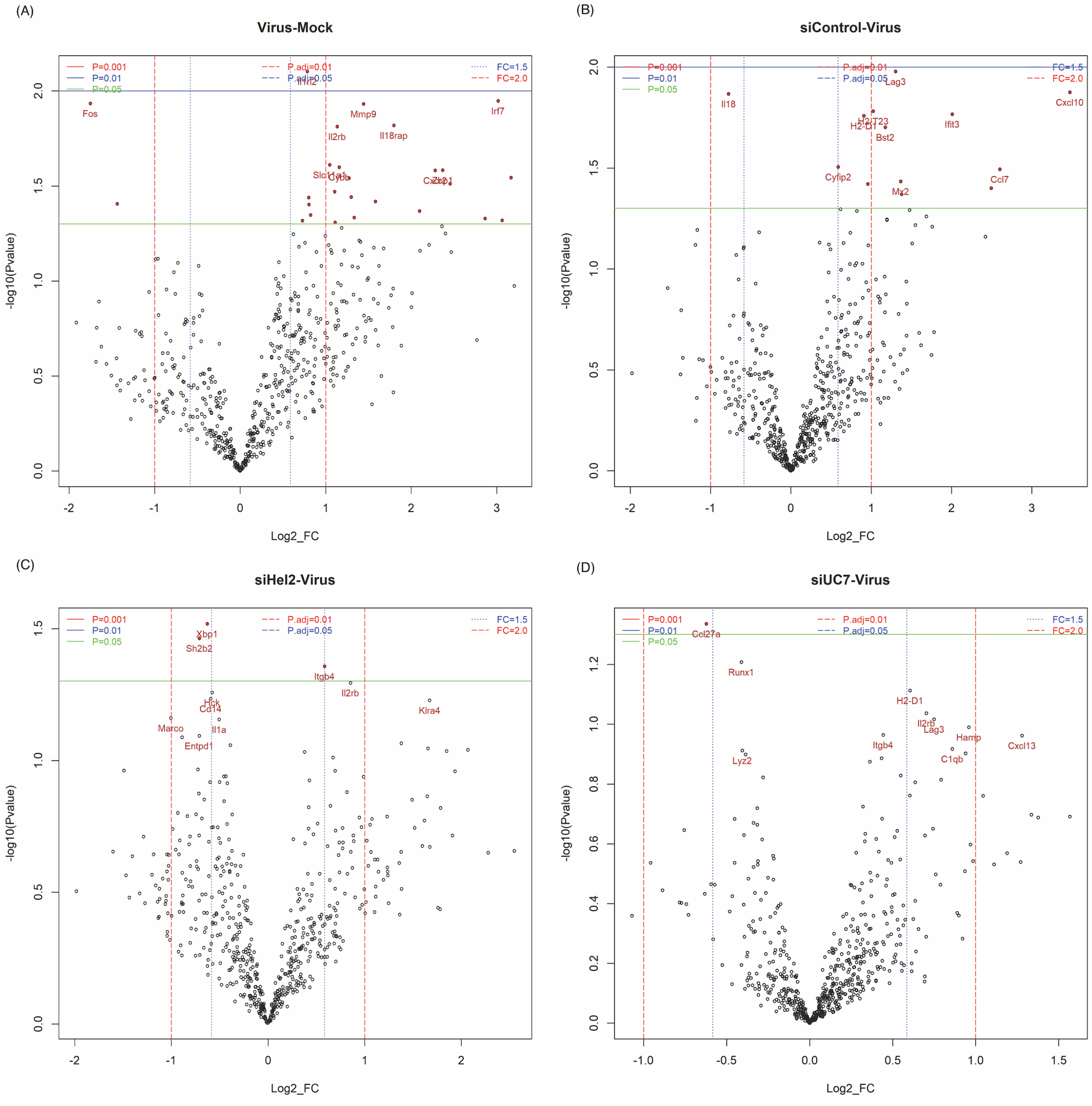


**Figure S6** *Volcano plots of differentially expressed immune genes in the lungs of K18-hACE2 female and male mice, 7–13-week-old, intranasally (IN) infected with either PBS (3), $10^4$ PFU/ 20 µL of SARS-CoV-2 or SARS-CoV-2 and siRNA treatment. (**A**) Virus alone vs. Mock, (**B**) siControl vs. Virus alone, (**C**) Virus alone vs. siHel2 treated and (**D**) Virus alone vs. siUC7 treated from day 6. Refer to Table S3 for raw data.*

Figure S7

(A)

| Formulation Name | DOTAP | MC3 | DSPC | Cholesterol | C16 PEG |
|---|---|---|---|---|---|
| DOTAP 40 | 40 | 10 | 10 | 37 | 3 |
| DOTAP 35 | 35 | 15 | 10 | 37 | 3 |
| DOTAP 30 | 30 | 20 | 10 | 37 | 3 |
| DOTAP 40C | 40 | 25 | 10 | 22 | 3 |

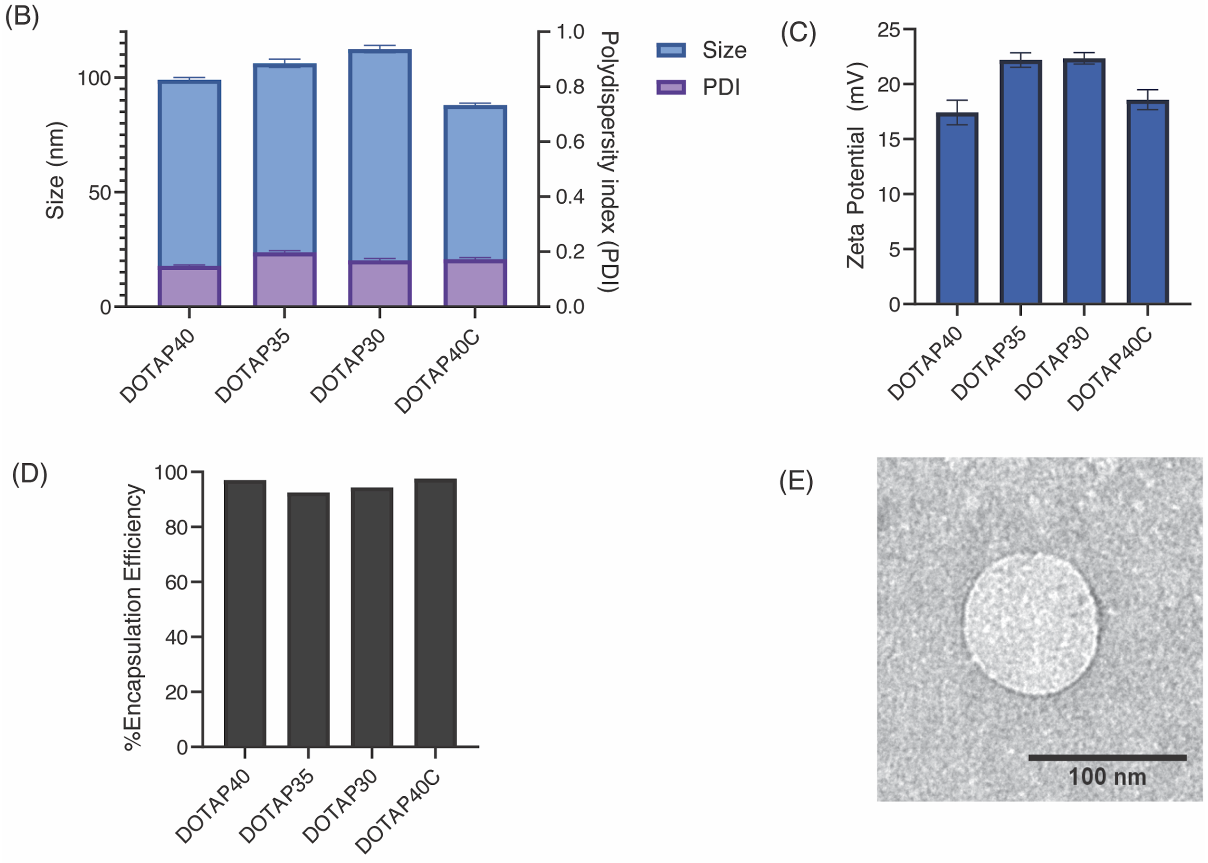


**Figure S7** *Optimization of reduced DOTAP nanoparticle formulations. (**A**) Composition of DOTAP nanoparticle formulations with Molar% of each lipid component reported. (**B**) Average size and polydispersity index of each DOTAP formulation was determined by dynamic light scattering (DLS). Error bars represent the S.E.M. of 5 runs (**C**) Average zeta potential reported in millivolts (mV) for each DOTAP formulation. Error bars represent the S.E.M. of 10 runs (**D**) SiRNA encapsulation efficiency of each formulation was determined using the Quant-IT Ribogreen assay. (**E**) Representative transmission electron microscope (TEM) image of DOTAP 40 nanoparticle formulation.*

Figure S8

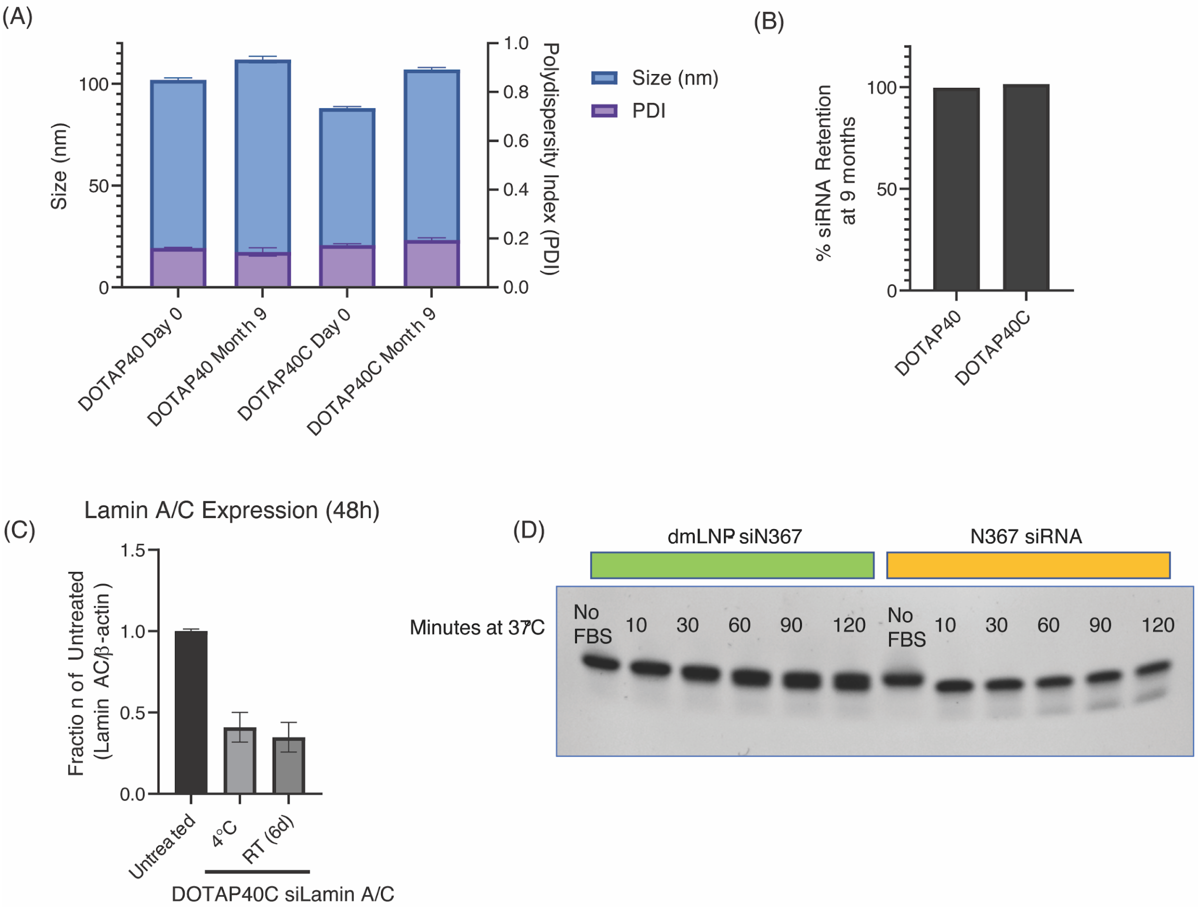


**Figure S8** *Supplemental Figure S8: Stability Evaluation of DOTAP40 and DOTAP40C nanoparticles. (**A**) Size and polydispersity measurements were carried out using dynamic light scattering (DLS) on DOTAP40 and DOTAP40C LNP formulations after synthesis and after 9 months of storage at 4°C. Error bars represent the S.E.M. of 5 runs. (**B**) Retention of siRNAs in LNPs after 9 months of storage. Ribogreen quantification of encapsulated siRNA in LNPs was evaluated at day 0 and again 9 months post synthesis. %siRNA retention= (final encapsulated siRNA concentration)/ (starting encapsulated siRNA concentration) *100. (**C**) DOTAP40C LNPs carrying Lamin A/C siRNA stored at either 4°C or room temperature for 6 days were dropped on NIH-3T3 cells at approximately 40nM and incubated for 48 hours. RNA was then extracted and 10ng of RNA was used in a Luna Universal One-Step RT-qPCR (New England BioLabs) to evaluate Lamin A/C expression. Error bars represent the S.E.M of triplicate wells. (**D**) Serum stability evaluation of dmLNP-siRNAs. 2.5ug of dmLNP-siN367 or N367 siRNA alone were incubated in 50ul of FBS (not heat inactivated) at 37°C for 0, 10, 30, 60, 90, and 120 minutes as done in (PMID: 19023647). RNase free water was added up to 200μL and RNA was subsequently extracted using phenol/chloroform (1:1 v/v) and centrifuged for at 14,000 rpm for 10 minutes at 4°C. The aqueous fraction was removed (approximately 25 μL), diluted 1/10, and electrophoresed on a non-denaturing 6% TBE polyacrylamide gel (Novex, Invitrogen) for 30 min at 200V. The gel was then stained with 2 μg/mL EtBr and images were acquired under 254 nm using an EZ Imager (Bio-Rad). Note: dmLNP is the updated name of the DOTAP40C formulation.*

**Table S1** *dsiRNA and siRNA sequences used in this study*

| siRNA | RNA sequence sense (5'-3') | RNA sequence antisense (5'-3') |
|---|---|---|
| **SiRdRp1** | rUrArUrGrGrGrUrUrGrGrGrArUrUrArUrCrCrUrArArArUGT | rArCrArUrUrUrArGrGrArUrArArUrCrCrCrArArCrCrCrArUrArArG |
| **SiRdRp1b** | rUrArUrGrGrGrUrUrGrGrGrArUrUrArCrCrCrUrArArArUGT | rArCrArUrUrUrArGrGrGrUrArArUrCrCrCrArArCrCrCrArUrArArG |
| **SiRdRp4** | rCrArArArGrArArUrArGrArGrCrUrCrGrCrArCrCrGrUrAGC | rGrCrUrArCrGrGrUrGrCrGrArGrCrUrCrUrArUrUrCrUrUrUrGrCrA |
| **SiHel1** | rUrGrUrUrGrArUrUrCrArUrCrArCrArGrGrGrCrUrCrArGAA | rUrUrCrUrGrArGrCrCrCrUrGrUrGrArUrGrArArUrCrArArCrArGrU |
| **SiHel2** | rArCrCrUrUrArUrArArUrUrCrArCrArGrArArUrGrCrUrGUA | rUrArCrArGrCrArUrUrCrUrGrUrGrArArUrUrArUrArArGrGrUrGrA |
| **SiUTR1** | rGrUrCrCrCrUrGrGrUrUrUrCrArArCrGrArGrArArArArCAC | rGrUrGrUrUrUrUrCrUrCrGrUrUrGrArArArCrCrArGrGrGrArCrArA |
| **SiUTR3** | rArUrArCrCrUrUrCrCrCrArGrGrUrArArCrArArArCrCrAAC | rGrUrUrGrGrUrUrUrGrUrUrArCrCrUrGrGrGrArArGrGrUrArUrArA |
| **siUC1** | rArArCrUrUrArUrGrUrArCrUrCrArUrUrCrGrUrUrU | rArCrGrArArUrGrArGrUrArCrArUrArArGrUrUrCrG |
| **siUC2** | rArCrUrUrArUrGrUrArCrUrCrArUrUrCrGrUrUrUrC | rArArCrGrArArUrGrArGrUrArCrArUrArArGrUrUrC |
| **siUC3** | rUrUrUrGrArArUrGrUrGrGrCrUrArArArUrCrUrGrA | rArGrArUrUrUrArGrCrCrArCrArUrUrCrArArArGrA |
| **siUC4** | rArCrCrArCrCrUrUrGrUrArGrGrUrUrUrGrUrUrArC | rArArCrArArArCrCrUrArCrArArGrGrUrGrGrUrUrC |
| **siUC5** | rUrGrGrArArCrCrArCrCrUrUrGrUrArGrGrUrUrUrG | rArArCrCrUrArCrArArGrGrUrGrGrUrUrCrCrArGrU |
| **siUC6** | rArArCrCrArCrCrUrUrGrUrArGrGrUrUrUrGrUrUrA | rArCrArArArCrCrUrArCrArArGrGrUrGrGrUrUrCrC |
| **siUC7** | rUrGrGrArCrUrGrArGrArCrUrGrArCrCrUrUrArCrU | rUrArArGrGrUrCrArGrUrCrUrCrArGrUrCrCrArArC |
| **siUC8** | rArCrUrGrGrArArCrCrArCrCrUrUrGrUrArGrGrUrU | rCrCrUrArCrArArGrGrUrGrGrUrUrCrCrArGrUrUrC |
| **siUC9** | rArUrGrArUrGrArUrUrArUrUrUrCrArArUrArArArA | rUrUrArUrUrGrArArArUrArArUrCrArUrCrArUrCrA |
| **siUC10** | rArCrUrArUrArUrGrUrUrArArArCrCrArGrGrUrGrG | rArCrCrUrGrGrUrUrUrArArCrArUrArUrArGrUrGrA |

| | | |
|---|---|---|
| **NC** | rCrArUrArUrUrGrCrGrCrGrUrArUrArGrUrCrGrCrG\rUrUAG | rCrUrArArCrGrCrGrArCrUrArUrArCrGrCrGrCrArArUrArUrGrGrU |
| **L362** | rGrArCrUrUrUrCrCrGrCrUrGrGrGrArCrUrUrUrC | rUrGrGrArArArGrUrCrCrCrCrArGrCrGrGrArArArG |
| **N367** | rCrUrGrArCrCrUrUrUrGrGrArUrGrGrUrGrCrUrUrC | rUrGrGrArArArGrUrCrCrCrCrArGrCrGrGrArArArG |
| **Lamin A/C** | rGrArCrUrUrGrGrUrGrUrGrGrArArGrGrCrGrCrArGrArACA | rUrGrUrUrCrUrGrCrGrCrCrUrUrCrCrArCrArCrCrArArGrUrCrArG |

*RNA bases denoted with r; DNA bases are capitalized*

**Table S2** *Normalised immune gene expression from the unsupervised hierarchical cluster analysis.*

**Table S3** *Normalised immune gene expression from the unsupervised hierarchical cluster analysis from day 6 volcano plot data in Figure S6.*

### Materials and Methods:

*Lipids/Reagents:*

The following lipids 1,2-dioleoyl-3-trimethylammonium-propane (DOTAP), 1,2-distearoyl-sn-glycero-3-phosphocholine (DSPC), Cholesterol, N-palmitoyl-sphingosine-1-{succinyl[methoxy(polyethylene glycol)2000]} (C16 PEG2000 Ceramide) were purchased from Avanti Polar Lipids (Alabaster, AL, USA). DLin-MC3-DMA was purchased from (MedChemExpress; Monmouth Junction, NJ, USA). All lipids were dissolved in ethanol and aliquoted in amber glass vials. Lipophilic dye DiL (DilC$_{18}$(3) or DiD (DilC$_{18}$(5) (1,1'-dioctadecyl-3,3,3',3'-tetramethylindocarbocyanine perchlorate) at 1mM stock in ethanol(Invitrogen; Carlsbad, CA, USA) was used to label nanoparticles at 0.5µM for biodistribution studies.

*dmLNP Synthesis:*

Lipids were prepared at a 40:25:10:22:3 (DOTAP:MC3:DSPC:Chol:PEG) molar ratio. Lipids in ethanol were mixed with nucleic acids in an aqueous phase at a mol cationic lipid: mol RNA (N:P) ratio of 3:1 using the NanoAssemblr Benchtop machine (Precision NanoSystems; Vancouver, BC, Canada). This machine contains a microfluidic chip by which the injected lipids and nucleic acids are mixed rapidly in a staggered herringbone pattern at a total flow rate of 12mLl/min. The controlled mixing of the aqueous and organic streams produces homogeneous nanoparticles. Immediately following the mixing process, the nanoparticles were diluted 1:4 with 1XPBS to reduce the amount of ethanol present in solution. The nanoparticle solution was further diluted with 1XPBS up to 15 mL and then concentrated using a 10 kDa Amicon ultra-15 filter (Millipore; Burlington, MA, USA) via centrifugation at 2,000 x G for 30 minutes. The flow through was discarded and another 15 mL 1XPBS was added to the column and centrifuged at 2,000 x G for 40 mins. The concentrated nanoparticles were then pushed through a 0.22µm filter and stored at 4°C.

*Characterization of dmLNPs:*

Nanoparticles were analyzed using dynamic light scattering (DLS) to determine polydispersity (PDI) and surface charge was determined by measuring zeta potential on a ZetaPals (Brookhaven Instruments Corporation; Holtsville, NY, USA). Concentration and relative nanoparticle size was determined using nanoparticle tracking analysis (NTA) on the Nanosight

(NS300) (Malvern Panalytical; Malvern, UK) and the qNANO Gold (IZON; Christchurch, New Zealand). For the NTA, samples were diluted 10,000XX in PBS and samples measured in triplicate technical replicates. A blue 488 laser was used to detect the LNPs, with a slide shutter level set to 1200X and the slider gain set to 146Y, and the syringe pump speed set to 30 using a flow-cell top plate module. For the qNano, a NP150 nanopore (iZON; Christchurch, New Zealand) was used to measure the LNPs. LNPs were diluted 40X in measuring solution (*6*) measured at 2 different pressures. Concentration was determined by measuring calibration beads at known concentrations and extrapolating particles/mL for each sample evaluated using the iZON control suite software (V3.4.2.48name, version).

To measure the amount of siRNA encapsulated inside the nanoparticles the Quant-IT Ribogreen assay was carried out (Molecular Probes; Eugene, OR, USA). The standard protocol was modified to include a 15 minute, 37°C incubation of the nanoparticles in the presence of 2% Triton to facilitate release of the encapsulated nucleic acids. %encapsulation = (siRNA-LNP in 2%Triton - siRNA-LNP in TE)/siRNA-LNP in 2%Triton-X100 (based on Leung 2012, PMID: 22962627).

*siRNAs:*

siRNAs or dsiRNAs targeted to SARS-CoV-2 were ordered from Integrated DNA Technologies (Coralville, IA, USA) as duplexed RNA. Target sequences for siRNAs are listed in Table S1. The negative control (NC) dsiRNA was purchased from IDT. Control siRNAs, N367 and L362 were designed towards the miRNA-N367 target site (*7-9*) and the 5'LTR (*10, 11*) of HIV-1 respectively, and were synthesized as duplexed RNA by IDT. Sequences for all dsiRNAs and siRNAs employed in this study are in supplementary Table 1. modUTR3 was designed as per supplementary Figure S2 and synthesized by IDT as an RNA duplex. HPLC purified and high quantity dsiRNAs and siRNAs, UTR3, Hel2, UC7, N367 were synthesized by the RNA/DNA Synthesis core at the City of Hope (Duarte, CA) and used in the *in vivo* experiments.

*Cells:*

HEK293 cells were purchased from ATCC (CRL-1573) and cultured in DMEM with 10% fetal bovine serum (FBS) at 37°C and 5% $CO_2$. Vero E6 cells were obtained from ATCC. Vero E6 and HEK293 cells were maintained in complete media; DMEM (Gibco-Invitrogen, Waltham, MA) supplemented with 10% heat inactivated foetal bovine serum (FBS) (30 min at 56°C, Gibco-Invitrogen, Waltham, MA) and 1% of antibiotic/glutamine preparation (100 U/ml penicillin G, 100 U/ml streptomycin sulphate, and 2.9 mg/ml of L-glutamine) (Gibco-Invitrogen, Waltham, MA). THP1-Dual™ cells (InvivoGen) were grown in RPMI 1640, 2 mM L-glutamine, 25 mM HEPES, 10% heat-inactivated FBS , 100 μg/ml Normocin™ and Pen-Strep (100 U/ml-100 μg/ml).

*Virus cultivation:*

SARS-CoV-2 VIC1 strain was obtained from the Peter Doherty Institute for Infection and Immunity and Melbourne Health, Victoria, Australia (*12*) and cultured in Vero E6 cells. Viral supernatant was concentrated in Amicon ® Ultra-15 Centrifugal Filter units (Merck, Germany) and viral titre determined by the viral immunoplaque assay as previously described (*13*).

*Cell transfection and siRNA Screening:*

Cells were seeded overnight in a 12-well plate to 70-80% confluency before transfecting siRNAs with either FuGENE 6 (Promega, Madison, WI) or Lipofectamine 2000 (Gibco-Invitrogen, Waltham, MA) in OptiMEM (Gibco-Invitrogen, Waltham, MA) as per manufacturer’s protocol. For the knockdown reporter assays, the CoV-2 ORF, helicase, 5’UTR) were cloned downstream of *Renilla* luciferase in a pSI-CHECK reporter vector and transfected with the siRNA into HEK293 in a 48-well plate. At 48 hrs post-transfection the levels of luciferase were measured and made relative to a control siRNA set at 100%. 250ng of pSI-CHECK plasmids and 0.75, 7.5 or 75 pmol of siRNA was used. Luciferase reporter activity was measured using a Dual-Luciferase® Reporter Assay System and measured on a GloMax Explorer microplate reader (Promega; Madison, WI, USA).

*Immunostimulation assay:*
THP1-Dual™ cell IRF and NFkB reporter gene expression were measured as per manufacturer's protocol (InvivoGen). NF-kB and IRF activation pathways were measured by assessing the activity of alkaline phosphatase and luciferase. 2’3’-cGAMP and LPS were obtained from InvivoGen (San Diego, CA).

*Negative staining electron microscopy:*
Nanoparticles diluted 1/100 in 1X PBS were absorbed to glow-discharged, carbon-coated 200 mesh EM grids. Samples were prepared by conventional negative staining with 1% (w/v) uranyl acetate. Electron microscopy images were taken on an FEI Tecnai 12 transmission electron microscope equipped with a Gatan OneView CMOS camera. Images were analyzed using ImageJ software (V1.52d).

*dmLNP-siRNA uptake evaluation in vivo:*
All animal experiments were approved by the City of Hope IACUC (20025). To determine if siRNA loaded nanoparticles show preferential lung accumulation we injected C57/BL6 mice IV with DiD-labeled nanoparticles at 1mg/kg siRNA dose. 24 hours after injection, mice were euthanized and the lung, liver, and spleen were removed. Organs were imaged for DiD fluorescence using a LagoX imager (Spectral Instruments Imaging, AZ) at an excitation and emission wavelength of 640 and 690 nm, respectively.

*Long-term stability:*
To determine the long-term stability of our LNPs we evaluated size and polydispersity using DLS and monitored siRNA encapsulation at 9 months post formulation.

*SARS-CoV-2 in vivo infection model:*
K18-hACE2 mice were purchased from the Jackson Laboratory (Bar Harbor, ME) and bred in-house at the Griffith University Animal Resource Center. Mice were intranasally infected with 1-4 x $10^4$ PFU (20 µL total volume) of live SARS-CoV-2 while under isoflurane anesthesia. Mice were subsequently treated with either LNP or DOTAP 40 complexed siRNAs, 5% sucrose (for LNP control) or PBS (for DOTAP 40 control) vehicle control (retro-orbitally) while under isoflurane anesthesia. Mice were monitored daily for weighing and clinical scoring. This work was conducted in a BSL3 approved animal facility at Griffith University (Animal ethics approval: MHIQ/07/20/AEC).

*Viral plaque and immunoplaque assay:*
For viral plaque assays, Vero E6 cells were infected with a MOI 0.002 of SARS-CoV-2 for 1hr before overlaying with 1% methylcellulose- viscosity 4,000 centipoise (Sigma- Aldrich, St. Louis, MO). Cells were incubated for 4 days at 37°C before fixing in 8% formaldehyde and stained with 1% crystal violet to visualize plaques. Viral immunoplaque assays were performed on Vero E6 cells as described previously (*13*) using recombinant monoclonal antibodies that

recognize SARS-CoV-2 (CR3022). Antibodies were obtained from Dr Naphak Modhiran and A/Prof. Dan Watterson (School of Chemistry and Molecular Biosciences, The University of Queensland, QLD, Australia). Virus titers were denoted as plaque forming units (PFU)/milliliter or PFU/grams of tissue.

*Viral copy number determination:*
To determine viral copy numbers in organ tissues, digital PCR against the N gene (CDC primers from IDT - SARS-CoV-2 N1) was performed in Quant-Studio 3D Digital PCR 20K chips (Thermo Scientific, Waltham, MA) ona ProFlex 2×Flat Block Thermal Cycler (Thermo Scientific, Waltham, MA). Results are analyzed on the QuantStudio 3D AnalysisSuite software (Thermo Scientific, Waltham, MA) and expressed as viral copies per gram of tissue.

*qRTPCR and gene expression analysis:* To evaluate the in vitro activity of LNPs carrying Lamin A/C siRNA, RT-qPCR analysis was carried out. RNA was isolated using the Maxwell RSC purification kit (Promega; Madison, WI, USA). A total of 10ng of RNA was used in Luna Universal One-Step RT-qPCR analysis (NEB; Ipswich, MA, USA) for Lamin A/C (5'-GAGAGGCTAAGAAGCAGC-3' and 5'-ACGCAGTTCCTCGCTGTAA-3'and β-actin 5'-GCTACAGCTTCACCACCACA-3' and 5'-TCTCCAGGGAGGAAGAGGAT-3'genes using the LightCycler96 real-time PCR system (Roche; Basel, Switzerland). Cycling conditions were as follows: reverse transcription (55°C for 10 min) and initial denaturation (95°C for 1 min) followed by 40 cycles of denaturation (95°C for 10 sec) and extension (60°C for 30 sec, with plate read). The fold change in gene expression was determined using the 2-DDCt method. The following qPCR primers were purchased from IDT.

*NanoString gene expression analysis:*
Immune gene expression analysis was undertaken using the NanoString nCounter analysis system (NanoString Technologies, Seattle, WA) and the commercially available nCounter Mouse PanCancer Immune Profiling panel kit. The PanCancer Immune profiling panel contains 730 genes of key inflammatory pathways 40 reference/housekeeping genes. The nCounter system directly detects and counts single-stranded nucleic acid via reporter probes affixed with fluorophore barcodes and biotinylated capture-probes attached to microscopic beads. Probes are then affixed to lanes in cartridges and read in a digital scanner. Following the manufacturer's protocol, 100 ng of total RNA extracted from tissue was hybridised with probes at 65 °C for 20 hours before being inserted into NanoString Prep Station where the target-probe complex was immobilised onto the analysis cartridge. Cartridges were read by the nCounter Digital Analyser for digital counting of molecular barcodes corresponding to each target at 555 fields of view.

*Nanostring Data analysis:*
Gene expression data was analysed using a combination of the Advanced Analysis Module in the nSolver™ Analysis Software version 4.0 from NanoString Technologies (NanoString Technologies, WA, USA), TIGR Multi-Experiment Viewer (http://mev.tm4.org) or the Limma package in the R Statistical Computing Environment. nSolver enables quality control (QC), normalisation, differential gene expression (DGE), Pathview Plots and immune cell profiling. Negative and positive controls included in probe sets were used for background thresholding, and normalizing samples for differences in hybridization or sample input respectively. Data was corrected for input volume via internal housekeeping genes using the geNorm algorithm. Genes that were expressed below 20 counts in more than 90% of samples were excluded from analysis. Differential gene expression between the treatment groups was determined using a

variance stabilised t-test. Pathway analysis was undertaken using the Kyoto Encyclopedia of Genes and Genomes (KEGG).

*Statistical analysis:*
All statistical analyses were performed using the statistical software package GraphPad Prism 9 and described in detail in respective figure legends.
